## Supplemental File S1 - Strain List for "Differential modulation of sensory response dynamics by cilia structure and intraflagellar transport within and across chemosensory neurons"

**Supplementary File 1.** List of strains used in this work.

| **Strain** | **Genotype** | **Source** |
| --- | --- | --- |
| PR802 | *osm-3(p802)* | CGC |
| PY12001 | *osm-3(oy156ts)* | This work |
| MX166 | *che-3(nx159ts)* | (Jensen et al., 2018) |
| VC513 | *grk-2(gk268)* | CGC |
| PY12006 | *osm-6(oy166[gfp_11_])* | This work |
| PY12014 | *kap-1(ok676)*; *osm-3(oy156ts)* | This work |
| PR811 | *osm-6(p811)* | CGC |
| PY12013 | *kap-1(ok676)*; *osm-3(p802)* | This work |
| PY1054 | *oyIs14[sra-6*p::*gfp]* | (Troemel et al., 1997) |
| PY11401 | *osm-6(p811)*; *oyEx[sra-6*p::*gfp]* | (Cornils et al., 2016) |
| PY12029 | *che-3(nx159ts)*; *oyIs14[sra-6*p::*gfp]* | This work |
| PY12030 | *kap-1(ok676)*; *osm-3(oy156ts); oyIs14[sra-6*p::*gfp]* | This work |
| PY12031 | *kap-1(ok676)*; *osm-3(p802)*; *oyIs14[sra-6*p::*gfp]* | This work |
| PY12005 | *kyIs602[sra-6*p::*GCaMP3]* | (Khan et al., 2022) |
| PY12032 | *kyIs602[sra-6*p::*GCaMP3]; kap-1(ok676)* | This work |
| PY12018 | *kyIs602[sra-6*p::*GCaMP3]; osm-3(p802)* | This work |
| PY12033 | *kyIs602[sra-6*p::*GCaMP3]; daf-10(e1387)* | This work |
| PY12034 | *kyIs602[sra-6*p::*GCaMP3]; osm-6(p811)* | This work |
| PY12019 | *kyIs602[sra-6*p::*GCaMP3]; kap-1(ok676)*; *osm-3(oy156ts)* | This work |
| PY12026 | *kyIs602[sra-6*p::*GCaMP3];che-3(nx159ts)* | This work |
| PY12035 | *osm-6(oy166[gfp_11_])*; *oyEx731[sra-6*p::*gfp_1-10_; sra-6*p::*myr-TagRFP]* | This work |
| PY12007 | *osm-6(oy166[gfp_11_])*; *oyEx682[sra-6*p::*gfp_1-10_]* | This work |
| PY12036 | *kap-1(ok676); osm-3(oy156ts); osm-6(oy166[gfp_11_])*; *oyEx682[sra-6*p::*gfp_1-10_]* | This work |
| PY12037 | *che-3(nx159ts); osm-6(oy166[gfp_11_])*; *oyEx682[sra-6*p::*gfp_1-10_]* | This work |
| PY6345 | *oyEx[sra-6*p::*osm-6*::*gfp]* | (Mukhopadhyay et al., 2007) |
| PY12023 | *osm-6(oy166[gfp_11_])*; *oyEx681[gpa-4Δ6*p::*gfp_1-10_; gpa-4Δ6*p::*mks-5*::*TagRFP]* | This work |
| PY12024 | *kap-1(ok676); osm-3(oy156ts); osm-6(oy166[gfp_11_]); oyEx681[gpa-4Δ6*p::*gfp_1-10_; gpa-4Δ6*p::*mks-5*::*TagRFP]* | This work |
| PY10210 | *oyIs88[gpa-4Δ6*p::*myr-gfp]* | From A. Maurya |
| PY12038 | *oyIs88[gpa-4Δ6*p::*myr-gfp]; osm-6(p811)* | From A. Maurya |
| PY12039 | *kap-1(ok676)*; *osm-3(p802)*; *oyIs88[gpa-4Δ6*p::*myr-gfp]* | This work |
| PY12028 | *kap-1(ok676)*; *osm-3(oy156ts)*; *oyIs88[gpa-4Δ6*p::*myr-gfp]* | This work |
| PY12027 | *grk-2(gk268)*; *oyIs88[gpa-4Δ6*p::*myr-gfp]* | This work |
| CX14887 | *kyIs598[gpa-6*p::*GCaMP2.2b]* | (Larsch et al., 2015) |
| PY12015 | *kyIs598[gpa-6*p::*GCaMP2.2b]*; *osm-6(p811)* | This work |
| PY12016 | *kyIs598[gpa-6*p::*GCaMP2.2b]*; *kap-1(ok676)*; *osm-3(p802)* | This work |
| PY12022 | *kyIs598[gpa-6*p::*GCaMP2.2b]*; *kap-1(ok676)*; *osm-3(oy156ts)* | This work |
| PY12017 | *kyIs598[gpa-6*p::*GCaMP2.2b]*; *grk-2(gk268)* | This work |
| PY12040 | *kyIs598[gpa-6*p::*GCaMP2.2b]*; *grk-2(gk268)*; *oyEx732[gpa-4Δ6*p::*grk-2*::*tagRFP]* | This work |
| PY12041 | *kyIs598[gpa-6*p::*GCaMP2.2b]*; *bbs-7(jhu590)* | This work |
| PY12042 | *kyIs598[gpa-6*p::*GCaMP2.2b]*; *bbs-7(ok1351)* | This work |
| PY12043 | *ebp-2(oy178[gfp_11_])*; *oyEx681[gpa-4Δ6*p::*gfp_1-10_; gpa-4Δ6*p::*mks-5*::*TagRFP]; oyEx733[gpa-4Δ6*p::*myr-TagRFP]* | This work |
| PY12044 | *ebp-2(oy178[gfp_11_])*; *osm-6(p811)*; *oyEx681[gpa-4Δ6*p::*gfp_1-10_; gpa-4Δ6*p::*mks-5*::*TagRFP]; oyEx733[gpa-4Δ6*p::*myr-TagRFP]* | This work |
| PY12045 | *oyEx734[gpa-4Δ6*p::*kap-1*::*gfp; gpa-4Δ6*p::*myr-TagRFP]* | This work |
| PY12046 | *osm-6(p811)*; *oyEx734[gpa-4Δ6*p::*kap-1*::*gfp; gpa-4Δ6*p::*myr-TagRFP]* | This work |
| PY12047 | *oyEx735[gpa-4Δ6*p::*osm-3*::*gfp; gpa-4Δ6*p::*myr-TagRFP]* | This work |
| PY12048 | *osm-6(p811)*; *oyEx735[gpa-4Δ6*p::*osm-3*::*gfp; gpa-4Δ6*p::*myr-TagRFP]* | This work |
| PY12049 | *oyEx736[gpa-4Δ6*p::*arl-13*::*TagRFP; gpa-4Δ6*p::*myr-gfp]* | This work |
| PY12050 | *osm-6(p811)*; *oyEx736[gpa-4Δ6*p::*arl-13*::*TagRFP; gpa-4Δ6*p::*myr-gfp]* | This work |
| PY12051 | *oyEx737[gpa-4Δ6*p::*odr-3*::*gfp; gpa-4Δ6*p::*myr-TagRFP]* | This work |
| PY12052 | *osm-6(p811)*; *oyEx737[gpa-4Δ6*p::*odr-3*::*gfp; gpa-4Δ6*p::*myr-TagRFP]* | This work |
| PY12053 | *oyEx738[gpa-4Δ6*p::*osm-9*::*gfp; gpa-4Δ6*p::*myr-TagRFP]* | This work |
| PY12054 | *osm-6(p811)*; *oyEx738[gpa-4Δ6*p::*osm-9*::*gfp; gpa-4Δ6*p::*myr-TagRFP]* | This work |
| PY12025 | *srx-64(oy193[gfp_11_])*; *oyEx681[gpa-4Δ6*p::*gfp_1-10_; gpa-4Δ6*p::*mks-5*::*TagRFP]* | This work |
| PY12055 | *srx-64(oy193[gfp_11_])*; *osm-6(p811)*; *oyEx681[gpa-4Δ6*p::*gfp_1-10_; gpa-4Δ6*p::*mks-5*::*TagRFP]* | This work |
| PY12056 | *osm-5(p813)*; *srx-64(oy193[gfp_11_])*; *oyEx681[gpa-4Δ6*p::*gfp_1-10_; gpa-4Δ6*p::*mks-5*::*TagRFP]* | This work |
| PY12057 | *kap-1(ok676)*; *osm-3(p802)*; *srx-64(oy193[gfp_11_])*; *oyEx681[gpa-4Δ6*p::*gfp_1-10_; gpa-4Δ6*p::*mks-5*::*TagRFP]* | This work |
| PY12058 | *bbs-7(jhu590)*; *srx-64(oy193[gfp_11_]); oyEx681[gpa-4Δ6*p::*gfp_1-10_; gpa-4Δ6*p::*mks-5*::*TagRFP]* | This work |
| PY12059 | *bbs-7(ok1351)*; *srx-64(oy193[gfp_11_])*; *oyEx681[gpa-4Δ6*p::*gfp_1-10_; gpa-4Δ6*p::*mks-5*::*TagRFP]* | This work |
| PY12004 | *odr-10(oy158[gfp_11_])*; *oyEx681[gpa-4Δ6*p::*gfp_1-10_; gpa-4Δ6*p::*mks-5*::*TagRFP]* | (Kyani-Rogers et al., 2022) |
| PY12060 | *odr-10(oy158[gfp_11_])*; *osm-6(p811)*; *oyEx681[gpa-4Δ6*p::*gfp_1-10_; gpa-4Δ6*p::*mks-5*::*TagRFP]* | This work |
| PY12061 | *odr-10(oy158[gfp_11_])*; *kap-1(ok676)*; *osm-3(p802)*; *oyEx681[gpa-4Δ6*p::*gfp_1-10_; gpa-4Δ6*p::*mks-5*::*TagRFP]* | This work |
| PY12062 | *odr-10(oy158[gfp_11_])*; *grk-2(gk268)*; *oyEx681[gpa-4Δ6*p::*gfp_1-10_; gpa-4Δ6*p::*mks-5*::*TagRFP]* | This work |
| PY12063 | *odr-10(oy158[gfp_11_])*; *grk-2(gk268)*; *oyEx681[gpa-4Δ6*p::*gfp_1-10_; gpa-4Δ6*p::*mks-5*::*TagRFP]; oyEx739[gpa-4Δ6*p::*grk-2*::*tagRFP]* | This work |
| PY12064 | *odr-10(oy194[FR])*; *oyEx681[gpa-4Δ6*p::*gfp_1-10_; gpa-4Δ6*p::*mks-5*::*TagRFP]* | This work |
| PY12065 | *odr-10(oy158[gfp_11_])*; *oyEx740[gpa-4Δ6*p::*gfp_1-10_; gpa-4Δ6*p::*myr-TagRFP]* | This work |
| PY12066 | *odr-10(oy158[gfp_11_])*; *kap-1(ok676)*; *osm-3(oy156ts)*; *oyEx740[gpa-4Δ6*p::*gfp_1-10_; gpa-4Δ6*p::*myr-TagRFP]* | This work |
| PY12067 | *odr-10(oy158[gfp_11_])*; *bbs-7(jhu590)*; *oyEx681[gpa-4Δ6*p::*gfp_1-10_; gpa-4Δ6*p::*mks-5*::*tagRFP]* | This work |
| PY12068 | *odr-10(oy158[gfp_11_])*; *bbs-7(ok1351)*; *oyEx681[gpa-4Δ6*p::*gfp_1-10_; gpa-4Δ6*p::*mks-5*::*tagRFP]* | This work |
| PY12069 | *srx-64(oy193[gfp_11_])*; *oyEx740[gpa-4Δ6*p::*gfp_1-10_; gpa-4Δ6*p::*myr-TagRFP]* | This work |
| PY12070 | *kap-1(ok676)*; *osm-3(oy156ts)*; *srx-64(oy193[gfp_11_])*; *oyEx740[gpa-4Δ6*p::*gfp_1-10_; gpa-4Δ6*p::*myr-TagRFP]* | This work |
| PY12071 | *oyIs88[gpa-4Δ6*p::*myr-gfp]*; *oyEx741[gpa-4Δ6*p::*grk-2*::*tagRFP]* | This work |
| PY12072 | *oyIs88[gpa-4Δ6*p::*myr-gfp]*; *osm-6(p811)*; *oyEx741[gpa-4Δ6*p::*grk-2*::*tagRFP]* | This work |
| PY12073 | *kap-1(ok676)*; *osm-3(oy156ts)*; *oyIs88[gpa-4Δ6*p::*myr-gfp]*; *oyEx741[gpa-4Δ6*p::*grk-2*::*tagRFP]* | This work |
| PY12074 | *oyEx742[odr-10*p::*srx-64*::*gfp; gpa-4Δ6*p::*myr-gfp]* | This work |
| PY12075 | *osm-6(p811)*; *oyEx742[odr-10*p::*srx-64*::*gfp; gpa-4Δ6*p::*myr-gfp]* | This work |
| PY12020 | *srx-64(oy195[SL2::gfp_11_])*; *oyEx681[gpa-4Δ6*p::*gfp_1-10_; gpa-4Δ6*p::*myr-TagRFP]* | This work |
| PY12021 | *osm-5(p813)*; *srx-64(oy195[SL2::gfp_11_])*; *oyEx681[gpa-4Δ6*p::*gfp_1-10_; gpa-4Δ6*p::*myr-TagRFP]* | This work |
| PY12080 | *kyIs598[gpa-6*p::*GCaMP2.2b]*; *osm-6(p811)*; *oyEx744[odr-10*p::*srx-64]* | This work |
| PY12076 | *odr-10(oy158[gfp_11_])*; *oyEx743[F16F9.3*p::*mCherry; gpa-4Δ6*p:: *gfp_1-10_]* | This work |
| PY12077 | *odr-10(oy158[gfp_11_])*; *kap-1(ok676)*; *osm-3(oy156ts)*; *oyEx743[F16F9.3*p::*mCherry; gpa-4Δ6*p:: *gfp_1-10_]* | This work |
| PY12078 | *srx-64(oy193[gfp_11_])*; *oyEx743[F16F9.3*p::*mCherry; gpa-4Δ6*p:: *gfp_1-10_]* | This work |
| PY12079 | *kap-1(ok676)*; *osm-3(oy156ts)*; *srx-64(oy193[gfp_11_])*; *oyEx743[F16F9.3*p::*mCherry; gpa-4Δ6*p:: *gfp_1-10_]* | This work |
