## Supplemental File S2 - Plasmid List for "Differential modulation of sensory response dynamics by cilia structure and intraflagellar transport within and across chemosensory neurons"

**Supplementary File 2.** List of plasmids used in this work.

| **Plasmid** | **Description** | **Source** |
| --- | --- | --- |
| PSAB1340 | 1kb_genomic*_osm-6_gfp_11__*1kb*_*genomic*_CRISPR* | This work |
| PSAB1341 | 1kb_genomic*_srx-64_gfp_11__1kb_*genomic*_CRISPR* | This work |
| PSAB1349 | 1kb_genomic*_srx-64_SL2*::*gfp_11__*1kb_genomic*_CRISPR* | This work |
| PSAB1280 | *gpa4Δ6*p::*gfp_1-10_* | (Kyani-Rogers et al., 2022) |
| PSAB1342 | *sra-6*p::*gfp_1-10_* | This work |
| PSAB1343 | *sra-6*p::*myr-TagRFP* | This work |
| PSAB1133 | *gpa4Δ6*p::*myr-TagRFP* | (Maurya et al., 2019) |
| PSAB1120 | *gpa4Δ6*p::*myr-GFP* | (Maurya et al., 2019) |
| PSAB1345 | *gpa4Δ6*p::*grk-2*::*TagRFP* | This work |
| PSAB1144 | *gpa4Δ6*p::*mks-5*::*TagRFP* | (Maurya et al., 2019) |
| PSAB1346 | *gpa4Δ6*p::*osm-9*::*gfp* | This work |
| PSAB1347 | *gpa4Δ6*p::*odr-3*::*gfp* | Ashish Maurya |
| PSAB1126 | *gpa4Δ6*p::*arl-13*::*TagRFP* | (Maurya et al., 2019) |
| PSAB1136 | *gpa4Δ6*p::*kap-1*::*gfp* | (Maurya et al., 2019) |
| PSAB1137 | *gpa4Δ6*p::*osm-3*::*gfp* | (Maurya et al., 2019) |
| PSAB1348 | *odr-10*p::*srx-64*::*gfp* | This work |
| PSAB1355 | *odr-10*p::*srx-64* | This work |
| PSAB1014 | *F16F9.3*p::*mCherry* | (Nechipurenko et al., 2016) |
