## Supplementary material for "Differential modulation of sensory response dynamics by cilia structure and intraflagellar transport within and across chemosensory neurons": Philbrook Supplemental

**SUPPLEMENTAL FIGURE LEGENDS**

**Figure S1. Characterization of the *osm-3(ts)* and endogenously tagged *osm-6* alleles.**

**A)** Sequence alignment of *C. elegans* kinesin motor subunits OSM-3, KLP-11, and KLP-20 with the *Chlamydomonas* kinesin motor subunits FLA10 and FLA8. The phenylalanine residue mutated to serine in FLA8-2 and engineered in OSM-3 to create *osm-3(oy156ts)* is indicated.

**B)** Percentage of animals of the indicated genotypes exhibiting defects in DiI uptake (Dyf) at 20°C or upon temperature upshift to 30°C for 4 hrs. n = 3 assays with 100 animals per genotype/condition.

**C)** ASH cilia length (left) or dye-filling (right) in animals carrying the *osm-6*::$gfp_{11}$ allele (*oy166*). Each dot is the value from a single neuron. Horizontal lines indicate the mean. Errors are SEM. ns: not significant. n = 3 assays, 100 animals per genotype (dye-filling).

**D)** Histogram of velocities of animals carrying the *osm-6*::$gfp_{11}$ *(oy166)* allele and $gfp_{1\text{-}10}$ expressed under the *sra-6* promoter in an extrachromosomal array, or expressing *sra-6*p::OSM-6::GFP from an extrachromosomal array. n ≥ 20 animals per genotype.

**Figure S2. Analysis of chemotaxis behaviors and odorant receptor expression in IFT mutants.**

**A,B)** Chemotaxis indices of animals of the indicated genotypes to the shown concentrations of odorants. Each dot is the chemotaxis index of a single assay, containing 100-300 adult hermaphrodites, n ≥ 5 assays performed over multiple days. ** and ***: different at P<0.01 and P<0.001 from corresponding wildtype (one-way ANOVA with Tukey's multiple comparisons test - A), ns: not significant. Dia – diacetyl; Pyr – pyrazine. (B) *srx-64*::$gfp_{11}$ is allele *srx-64(oy193)*, $gfp_{1\text{-}10}$ is expressed under the *gpa-4Δ6* promoter.

**C)** Representative image of endogenously tagged SRX-64::split-GFP expression in AWA cilia in wildtype or *osm-5(p813)* mutants. Numbers at bottom right indicate the percentage of animals exhibiting the shown phenotype; n ≥ 20. White/yellow arrowheads indicate cilia tip/base. Scale bar: 5 µm.

**D)** Representative images (top) and quantification (bottom) of SRX-64::SL2::split-GFP fluorescence in the AWA soma in the indicated genotypes. Scale bar: 5 µm. Each dot is the value from a single AWA neuron. ***: different at P<0.001 from wildtype (t-test).

Horizontal lines in all plots indicate mean. Errors are SEM. Data shown are from a minimum of 2-3 independent experiments.

**Figure S3. A subset of signaling molecules is localized to the PCMC branches in AWA.**

**A)** Representative images of AWA sensory endings in *kap-1; osm-3(ts)* mutants prior to and following temperature shift to the restrictive temperature for 24 hrs. Numbers at bottom right indicate the percentage of animals exhibiting PCMC branches; n ≥ 26.

**B-D)** Representative images of indicated reporter-tagged proteins in wildtype and *osm-6(p811)* mutants. Numbers at bottom right indicate the percentage of neurons exhibiting the shown phenotype; n ≥ 20 each.

White/yellow arrows indicate cilia tip/base. Arrows indicate PCMC branches. Scale bars: 5 µm.

**Figure S4. *grk-2* shapes odorant response amplitudes in AWA.**

**A)** Quantification of peak fluorescence intensity changes for data shown in Figure 4A,B. * and ***: different at P<0.05 and P<0.001 from indicated genotype (one-way ANOVA with Tukey's multiple comparisons test - left, t-test – right). ns: not significant.

**B)** Representative images of AWA cilia in wildtype and *grk-2(gk268)* animals expressing *gpa-4Δ6*p::*myr-GFP*. White/yellow arrows indicate cilia tip/base. Numbers at top right indicate percentage of animals exhibiting the shown phenotype; n ≥ 20 each. Scale bar: 5 μm.

**Figure S5. Diacetyl adaptation in AWA requires the GRK-2 kinase and IFT.**

**A)** Adaptation of peak AWA fluorescence upon repeated pulses of diacetyl in the indicated genotypes, normalized to the maximum peak $\Delta F/F_0$. n ≥ 14 animals for each data point. ** and ***: different at P<0.01 and P<0.001 from wildtype (second way ANOVA with repeated measures).

**B)** Percentage recovery of ODR-10::split-GFP fluorescence 40 secs after photobleaching the AWA ciliary stalk (red) or a similarly sized region of PCMC branches (black) in *osm-6(p802)* mutants. Data in red are also shown in Figure 5B. **: different at P<0.01 from indicated (t-test).

**C)** Representative images (left) and quantification (right) of ODR-10::split-GFP levels in wildtype and *grk-2(gk268)* animals. Wildtype *grk-2* sequences tagged with tagRFP were expressed in AWA under the *gpa-4Δ6* promoter. ** and ***: different at P<0.01 and P<0.001 between indicated values (one-way ANOVA with Tukey's multiple comparisons test).

**D)** A predicted BBSome-binding motif ([W/F/Y]R) is located within helix 8 of ciliary GPCRs (Yang et al., 2020) but absent in SRX-64. Conserved BBSome-binding residues are shown in bold.

**E)** Representative images of animals expressing wildtype ODR-10::split-GFP or a mutant protein containing a FR-to-AA substitution in the predicted BBSome binding site shown in D at the endogenous locus. Numbers at bottom right indicate percentage of animals exhibiting the shown phenotype; n ≥ 20 each.

**F)** Adaptation of peak AWA fluorescence upon repeated pulses of diacetyl in the indicated genotypes, normalized to the maximum peak $\Delta F/F_0$. $n \geq 9$ animals for each data point. *,** and ***: different at P<0.05, P<0.01, and P<0.001 from wildtype (second way ANOVA with repeated measures).

**G)** Summary of hypothesized IFT-dependent and independent removal of ODR-10 and SRX-64 GPCRs, respectively, during odorant adaptation. Upon interaction with diacetyl, ODR-10 is phosphorylated by GRK-2 and removed from cilia via BBSome and IFT. ODR-10 is then subsequently trafficked retrogradely to the AWA soma. In contrast, upon pyrazine-mediated activation of SRX-64, this receptor may be removed from cilia via ectocytosis.

In all images, yellow/white arrowheads indicate cilia base/cilia tip. Each dot in scatter plots is the measurement from a single AWA neuron. Horizontal lines indicate mean. Errors are SEM. Data shown are from a minimum of 2-3 independent experiments. ns: not significant. Scale bars: 5 μm.

**Movie S1.** Movement of endogenously tagged OSM-6::split-GFP within the primary AWA ciliary stalk in wildtype animals. Arrowhead marks cilia base, distal end of cilia is at top. Images were captured at 4 Hz and played back at 20 frames per sec.

**Movie S2.** Movement of endogenously tagged EBP-2::split-GFP movement within dendritic branches in *osm-6(p811)* mutants. Arrowhead marks PCMC branches, distal end of cilia is at top. Images were captured at 4 Hz and played back at 20 frames per sec.

**Movie S3.** Trafficking of endogenously tagged ODR-10::split-GFP puncta in the AWA dendrite upon addition of $10^{-4}$ dilution of diacetyl. Cilia at left, soma at right. Images were captured at 4 Hz and played back at 20 frames per sec.

**Movie S4.** Trafficking of endogenously tagged SRX-64::split-GFP puncta in the AWA dendrite upon addition of $10^{-4}$ dilution of pyrazine. Cilia at left, soma at right. Images were captured at 4 Hz and played back at 20 frames per sec.

**S1 Table.** Quantification of OSM-6::GFP or OSM-6::split-GFP anterograde movement in ASH cilia.

| **Strain: Genotype** | **Condition[3]** | **Average velocity (µm/sec ± SEM)** | **Average number of IFT events per 30s (± SEM)** | **Number of animals imaged** |
|---|---|---|---|---|
| PSAB6345: Ex[*sra-6*p::*osm-6::gfp*] | 20°C | 0.79 ± 0.01 | 15.5 ± 1.36 | 18 |
| PSAB12007: *osm-6(oy166)*[1]; *oyEx682*[2] | 20°C | 0.81 ± 0.01 | 15.7 ± 0.93 | 24 |
| PSAB12007: *osm-6(oy166)*[1]; *oyEx682*[2] | 20°C | 0.95 ± 0.01 | 23.1 ± 0.56 | 20 |
| PSAB12007: *osm-6(oy166)*[1]; *oyEx682*[2] | 1 hr 27°C | 0.89 ± 0.01 | 20.5 ± 0.75 | 21 |
| PSAB12007: *osm-6(oy166)*[1]; *oyEx682*[2] | 1 hr 30°C | 1.01 ± 0.01 | 23.1 ± 0.72 | 22 |
| PSAB12036: *kap-1(ok676); osm-3(oy156ts); osm-6(oy166)*[1]; *oyEx682*[2] | 20°C | 0.78 ± 0.01 | 19.1 ± 0.58 | 24 |
| PSAB12036: *kap-1(ok676); osm-3(oy156ts); osm-6(oy166)*[1]; *oyEx682*[2] | 1 hr 27°C | 0.80 ± 0.01 | 15.4 ± 0.93 | 24 |
| PSAB12036: *kap-1(ok676); osm-3(oy156ts); osm-6(oy166)*[1]; *oyEx682*[2] | 1 hr 30°C | 0.70 ± 0.02 | 3.86 ± 0.81 | 21 |
| PSAB12007: *osm-6(oy166)*[1]; *oyEx682*[2] | 15°C | 0.92 ± 0.01 | 20.6 ± 0.77 | 20 |
| PSAB12007: *osm-6(oy166)*[1]; *oyEx682*[2] | 3 hr 25°C | 0.86 ± 0.01 | 20.4 ± 0.76 | 22 |
| PSAB12037: *che-3(nx159ts); osm-6(oy166)*[1]; *oyEx682*[2] | 15°C | 0.84 ± 0.01 | 17.6 ± 0.80 | 20 |
| PSAB12037: *che-3(nx159ts); osm-6(oy166)*[1]; *oyEx682*[2] | 3 hr 25°C | 0.84 ± 0.01 | 9.55 ± 1.30 | 20 |

[1]*osm-6(oy166): osm-6*::$gfp_{11}$

[2]*oyEx682: Ex[sra-6*p::$gfp_{1-10}$*]*

[3]*kap-1; osm-3(ts)* and associated control animals were grown at 20°C prior to the indicated temperature upshifts. *che-3(ts)* and associated control animals were grown at 15°C prior to the indicated temperature upshifts.

**S2 Table.** Quantification of OSM-6::split-GFP anterograde movement in the AWA cilia stalk.

| Strain: Genotype | Condition[3] | Average velocity (µm/sec ± SEM) | Average number of IFT events per 30s (± SEM) | Number of animals imaged |
|---|---|---|---|---|
| PSAB12023: *osm-6(oy166)*[1]*; oyEx681*[2] | 20°C | 0.60 ± 0.01 | 10.2 ± 1.58 | 22 |
| PSAB12023: *osm-6(oy166)*[1]*; oyEx681*[2] | 1.5 hr 30°C | 0.64 ± 0.01 | 11.7 ± 1.71 | 18 |
| PSAB12024: *kap-1(ok676); osm-3(oy156ts); osm-6(oy166)*[1]*; oyEx681*[2] | 20°C | 0.79 ± 0.02 | 5.24 ± 1.35 | 21 |
| PSAB12024: *kap-1(ok676); osm-3(oy156ts); osm-6(oy166)*[1]*; oyEx681*[2] | 1.5 hr 30°C | 0.60 ± 0.05 | 0.29 ± 0.29 | 21 |

[1]*osm-6(oy166): osm-6*::*gfp*$_{11}$

[2]*oyEx681: Ex[gpa-4Δ6*p::*gfp*$_{1-10}$*]*

[3]Animals were grown at 20°C prior to the indicated temperature upshifts.

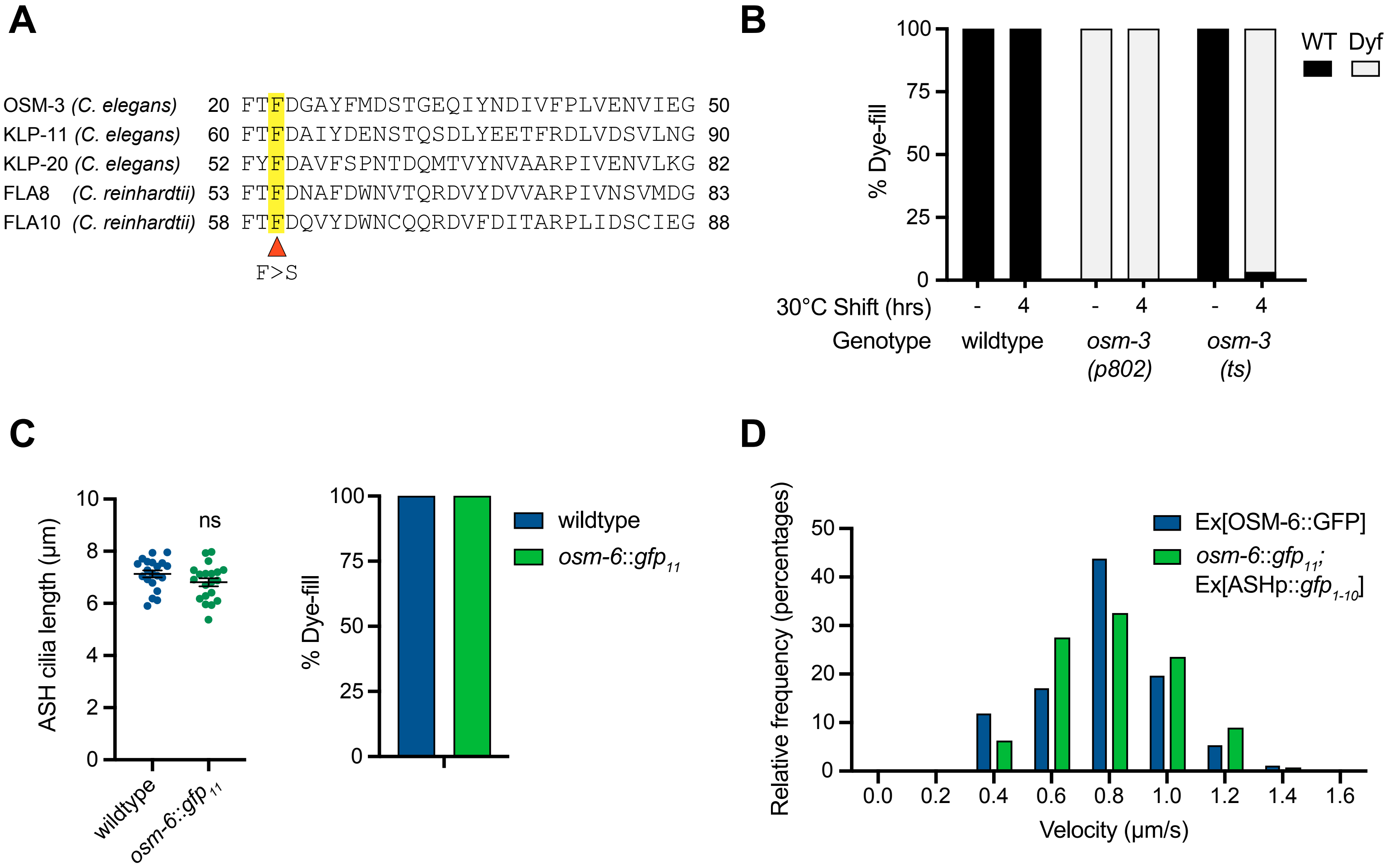

**Philbrook et al., Figure S1**

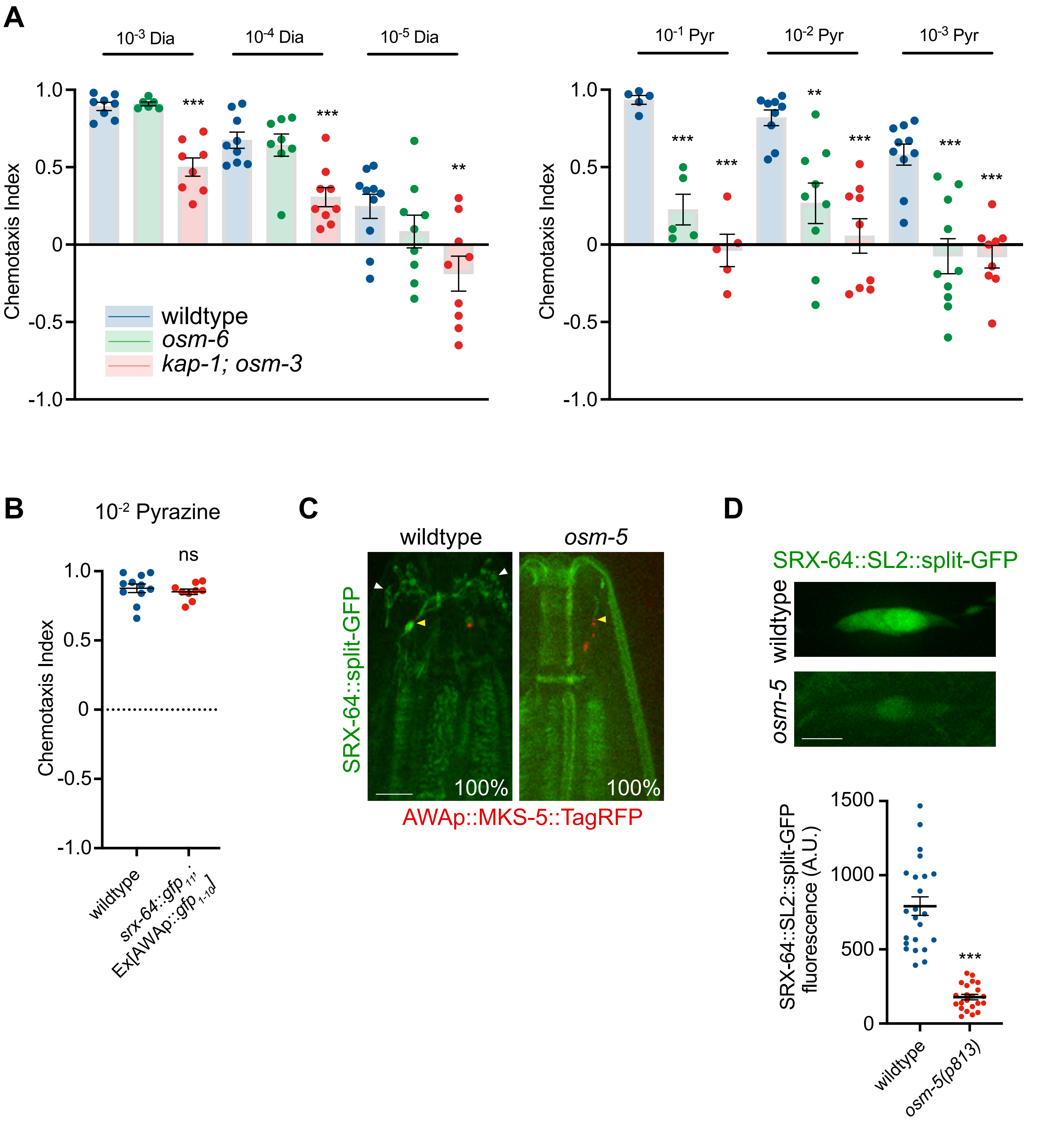

Philbrook et al., Figure S2

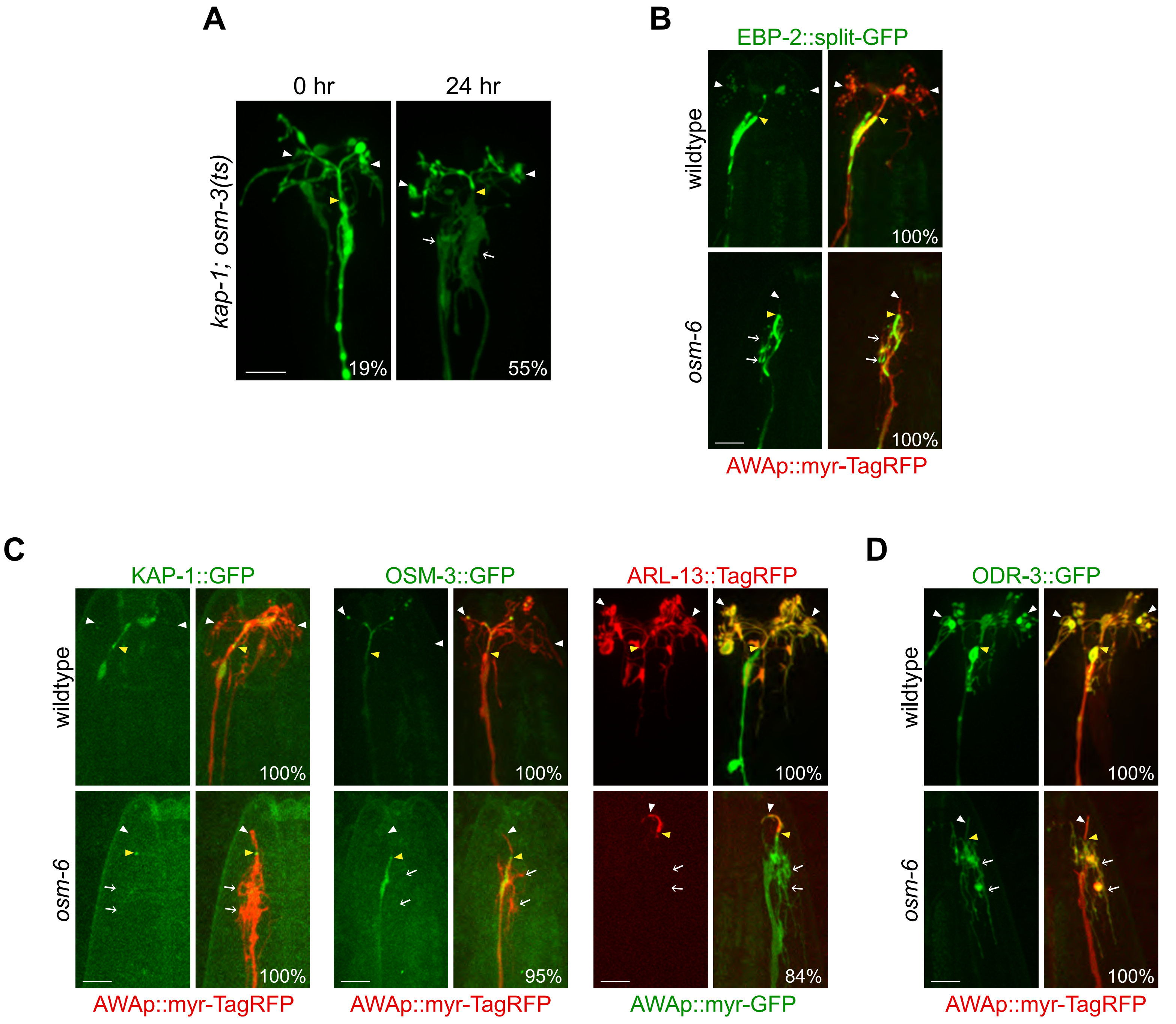

**Philbrook et al., Figure S3**

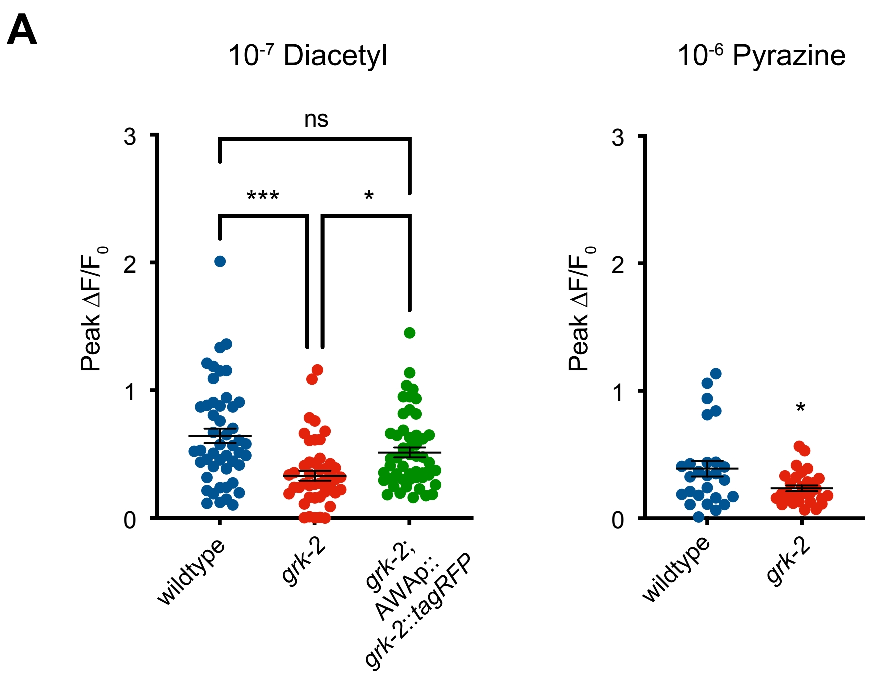

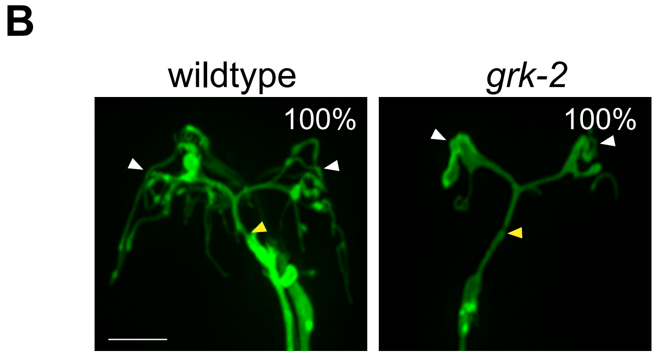

Philbrook et al., Figure S4

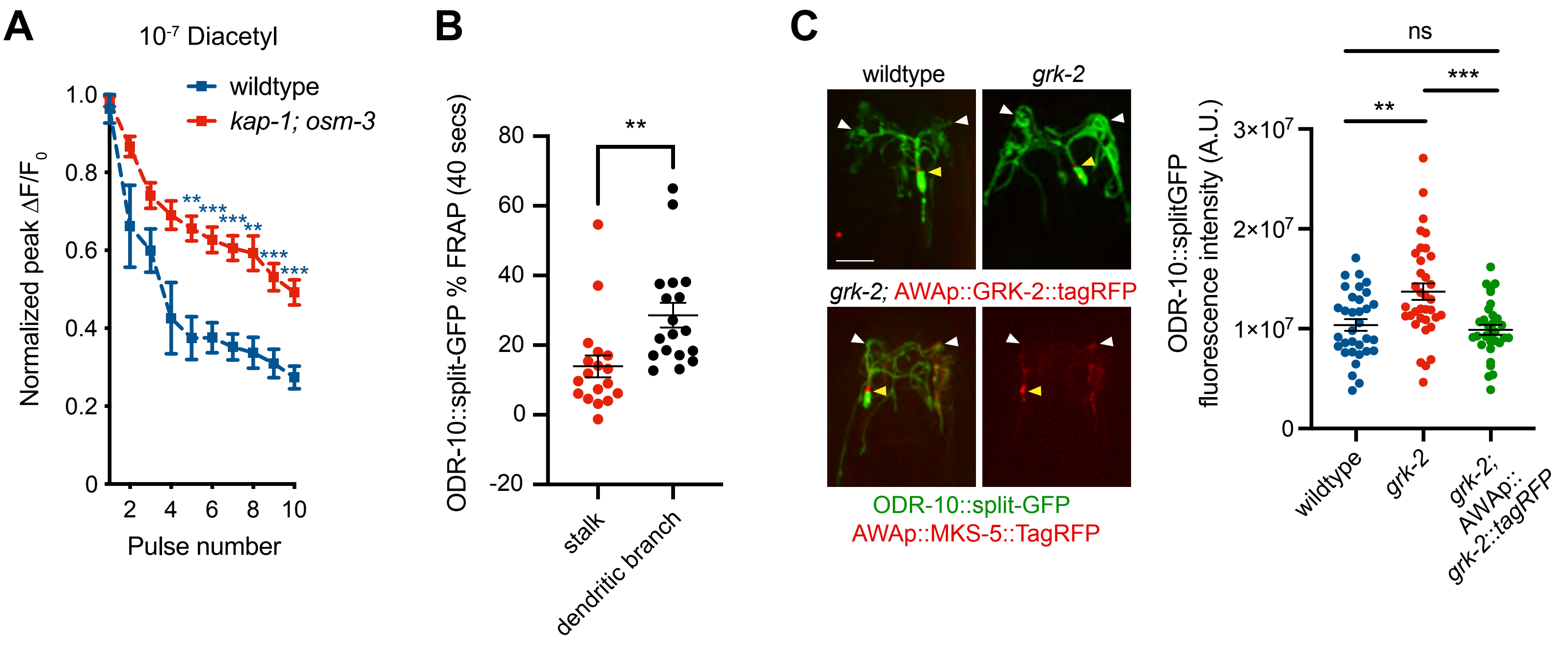

D

| ciliary GPCR | Helix 8 |
|---|---|
| *CCKAR* | NKR**FR**LGFMATF |
| *DRD1* | FNAD**FR**KAFSTLLGC |
| *DRD2S* | NIE**FR**KAFLKILHC |
| *FFAR4* | LCRNE**WK**KIFCC |
| *GALR2* | SKH**FR**KG**FR**TICA |
| *GALR3* | SRH**FR**AR**FR**RLWPC |
| *GPR19* | NAN**FR**RGMKETF |
| *GPR83* | NEN**FR**IELKALL |
| *GPR88* | T**WR**NEE**FR**RSVRSV |
| *HTR6* | MRD**FK**RALGRFVPC |
| *KISS1R* | GSHFRQA**FR**RVCPC |
| *MC4R* | SQELRKT**FK**EIIC |
| *MCH1R* | CET**FR**KRLVLSVK |
| *NMUR1* | SSR**FR**ETFQEALCLGACC |
| *NPFFR1* | NEN**FR**RGFQAA**FR**ARLC |
| *P2RY1* | GDT**FR**RRLSRATR |
| *PRLHR* | HDS**FR**EELRKLL |
| *Pgr15L* | NRS**FR**AKLRSISS**FR**M |
| *NPY2R* | NSN**YR**KAFLSA**FR**C |
| *QRFPR* | EN**FK**KNVLSAVC |
| *SSTR3* | S**YR**FKQG**FR**RILL |
| *TGR5* | DQRYTAP**WR**AAAQ |
| *Ce:* ODR-10 | RD**FR**RTIFNFLC |
| *Ce:* SRX-64 | NPEIRKFLLGKK |

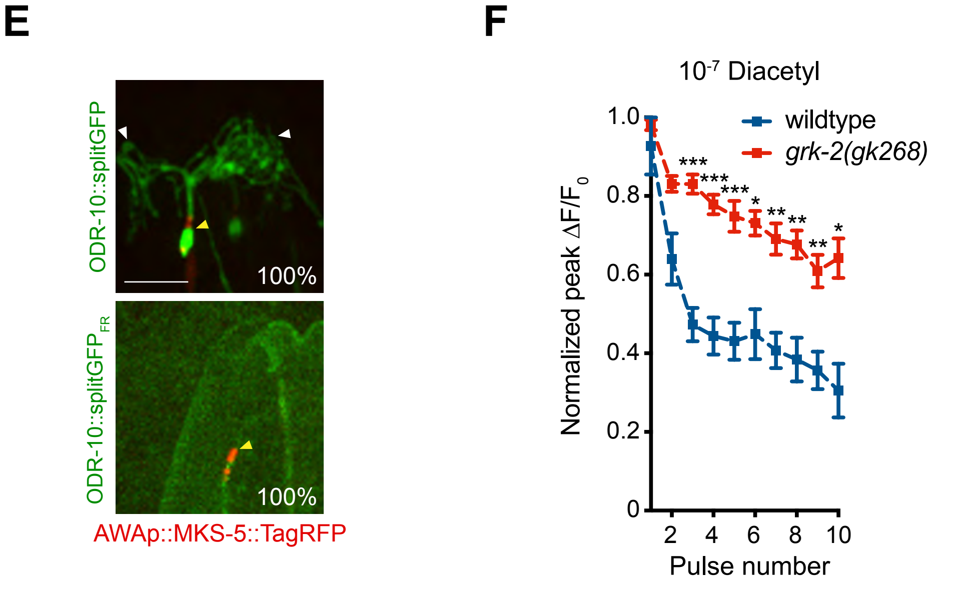

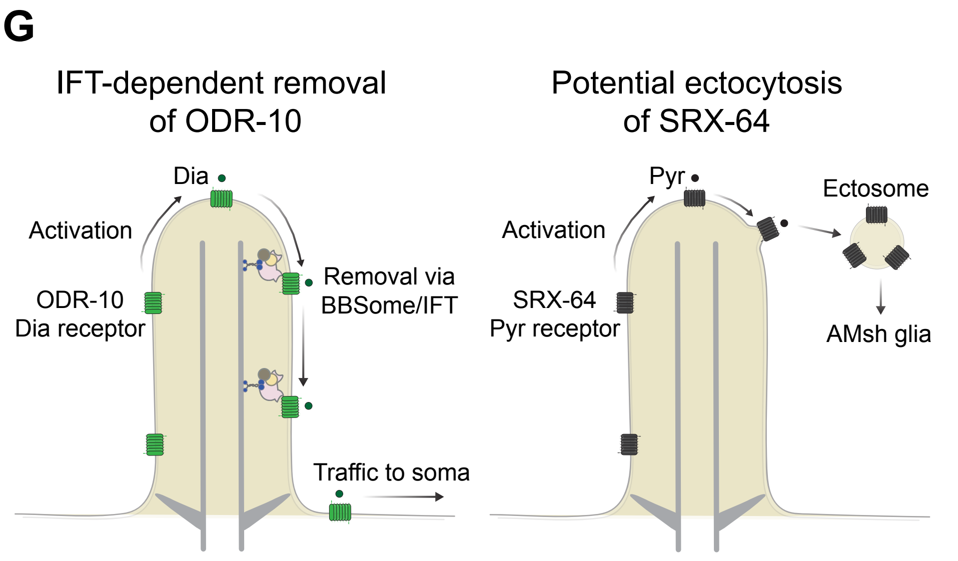

**Philbrook et al., Figure S5**
